# A machine learning approach to identifying objective biomarkers of anxiety and stress

**DOI:** 10.1101/745315

**Authors:** Arjun Ramakrishnan, Adam Pardes, William Lynch, Christopher Molaro, Michael Louis Platt

**Author notes:** ***Corresponding author***, Arjun Ramakrishnan, Department of Neuroscience, University of Pennsylvania, Philadelphia, PA 19104.

## Abstract

Anxiety and stress-related disorders are highly prevalent and debilitating conditions that impose an enormous burden on society. Sensitive measurements that can enable early diagnosis could mitigate suffering and potentially prevent onset of these conditions. Self-reports, however, are intrusive and vulnerable to biases that can conceal the true internal state. Physiological responses, on the other hand, manifest spontaneously and can be monitored continuously, providing potential objective biomarkers for anxiety and stress. Recent studies have shown that algorithms trained on physiological measurements can predict stress states with high accuracy. Whether these predictive algorithms generalize to untested situations and participants, however, remains unclear. Further, whether biomarkers of momentary stress indicate trait anxiety – a vulnerability foreshadowing development of anxiety and mood disorders – remains unknown. To address these gaps, we monitored skin conductance, heart rate, heart rate variability and EEG in 39 participants experiencing physical and social stress and compared these measures to non-stressful periods of talking, rest, and playing a simple video game. Self-report measures were obtained periodically throughout the experiment. A support vector machine trained on physiological measurements identified stress conditions with ~96% accuracy. A decision tree that optimally combined physiological and self-report measures identified individuals with high trait anxiety with ~84% accuracy. Individuals with high trait anxiety also displayed high baseline state anxiety but a muted physiological response to acute stressors. Overall, these results demonstrate the potential for using machine learning tools to identify objective biomarkers useful for diagnosing and monitoring mental health conditions like anxiety and depression.

## Introduction

Anxiety and other stress-related disorders are the most common forms of mental illness with over 250 million affected people worldwide (1). Thirty-one percent of U.S. adults will experience an anxiety disorder at some time in their lives (2–4) and 40% of graduate students may suffer from it at some point (5). These conditions are highly treatable and early intervention improves patient outcomes (6). Unfortunately, sensitive and reliable biomarkers to enable early diagnosis and personalized intervention strategies remain elusive (7).

Anxiety disorders are defined by excessive or uncontrolled anticipation of uncertain threat that can lead people to seek medical attention (8). Various social and biological factors (9) may render some individuals more vulnerable to developing anxiety disorders (10). Trait anxiety is operationalized as the proneness to experience maladaptive anxious states and foreshadows vulnerability to developing a true anxiety disorder (11). Stress can be a major anxiogenic process. Stressful events induce physiological responses, or allostasis, (12) that generate a transient increase in state anxiety (11), and early life stressful events or chronic, unmanaged stress may predispose individuals to developing anxiety disorders (9). Understanding the relationships between stress response and state and trait anxiety may help unlock the mechanisms underlying development of anxiety disorders.

Currently available screening tools to monitor stress response, like the VAS and SUDS, are quick and reliable but are overtly interventional, not amenable to continuous monitoring, vulnerable to biased reporting, and suffer from diminished sensitivity over repeated assessments (13–17). To mitigate biased reporting, studies have begun to examine the efficacy of biochemical and physiological measures, which manifest spontaneously and are harder to conceal, to supplement self-reported measures of stress. While biochemical assays are invasive, expensive, not amenable to continuous measurements, and vulnerable to diurnal cycles(18–21), electrophysiological measures are non-invasive and more suitable for continuous monitoring. Measures like galvanic skin response (GSR) (22–29), heart rate (HR)(24–26,28,30), heart rate variability (HRV)(27,31,32), and electroencephalography (EEG) (33–36) are increasingly being used to assess stress response. The relationship between these physiological measures and stress response, and both state and trait anxiety, however, remains poorly understood (37).

Several recent studies have used new developments in machine learning to integrate features from more than one physiological measure to evaluate stress response with high accuracy (24,25,29,33,38–42). These algorithms, however, have not always been validated in conditions outside the ones in which they were trained. Thus, it remains unclear whether measures obtained and algorithms developed in one condition can generalize to an untested condition. Similarly, whether the algorithms trained on physiological responses to one stressor can predict physiological responses to another stressor is not known. Finally, it is unclear whether algorithms trained to predict stress response in one cohort of individuals generalize to new cohorts of individuals.

To address these questions, we studied participants who were subjected to two types of stress tests, cold pressor test (CPT) and the trier social stress test (TSST)(43,44). During the session, participants also engaged in non-stressful activities that involved talking, rest, and playing a simple video game. Both standardized self-reports and physiological data were collected prior to and during the session. We first assessed the efficacy of different physiological measures to infer epochs in which stressors were applied. We then developed a novel algorithm trained on the best features from these different physiological measurements to identify a stressful epoch with ~96% accuracy. Next, by combining the algorithm-based predictions with self-report measures, we developed a procedure to identify individuals with high self-reported trait anxiety with ~84% accuracy. Finally, we examined the relationship between trait anxiety, state anxiety and the physiological response to stress and found that individuals with high trait anxiety also displayed high baseline state anxiety but a muted physiological response to acute stressors.

## Results

In this study, 39 participants experienced two types of stressors, cold pressor test (CPT) and Trier social stress test (TSST), in the following order (Fig 1). They also played a simple decision making game both before CPT and after TSST. They were engaged in casual conversation (Talk epoch) and also received a rest period during the session (Fig 1). Overall, a session lasted approximately 60 minutes (refer to Methods for more details).

**Figure 1:**
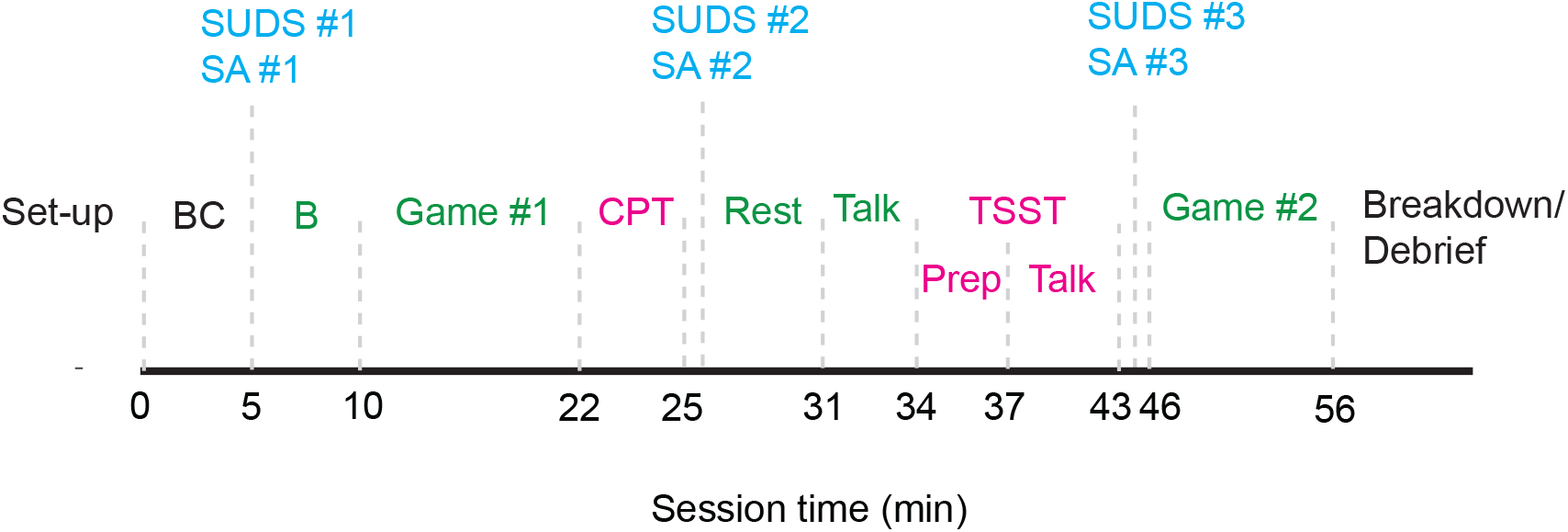
Study protocol. Following initial briefing and set-up, participants were asked to relax for 10 minutes. The first 5 minutes were considered baseline control (BC) and the physiological readings during the session were measured relative to this epoch. The next 5 minutes were considered baseline period (B). Participants’ self-reports – both subjective units of distress (SUDS) and state anxiety (SA) -- were obtained during the baseline period. Participants then played a simple decision making game (Game #1) for 10 minutes after 2 minutes of practice. Following this they were subjected to the Cold Pressor Test (CPT). After CPT, a 6 minute relaxation period began. During this time, the second self-report measurements of state anxiety were obtained. This was followed by 3 minute ‘talk’ epoch. This was followed by TSST: 3 min for speech preparation (Prep), 5 min for the speech (‘talk’) followed by a minute for mental countdown. After TSST, self report surveys were obtained for the third and final time. Participants then played the same video game again (‘Game #2’). After this, they were debriefed and the equipment was removed. Stress induction epochs are shaded magenta, non-stressful epochs are shaded green.

### Self-reported responses to induced stress

To determine each participants’ stress response, we first examined their self-report scores for subjective units of distress (SUDs) and state anxiety (SA). An example participant (P25) whose data is plotted in Fig 2A reported a strong response to the applied stressors. This participant showed a low baseline score (SUDs = 10; SA = 28), a slightly elevated score post-CPT (SUDs = 15; SA = 34) and a much more elevated score post-TSST (SUDs = 60; SA = 65). Some participants, like the one whose data is plotted in Fig 2B (P13), did not report an increase in stress. For this participant, scores remained close to baseline for SA (Baseline: 25; post-CPT: 26; post-TSST: 24) and decreased from baseline for SUDS (Baseline: 60; post-CPT: 40; post-TSST: 40). We chose an arbitrary threshold for stress response (Change in SUDS > 50% and SA > 30%) to identify self-reported strong responders from weak/non-responders for illustrative purposes (Fig 2 C-D). Twenty-three of 39 participants qualified as strong responders based the arbitrary threshold (red dots in Fig 2C-D). For the population, the average change in state anxiety was 90±9% (Fig. 2E) and the change in SUDS scores was 320±61% (Fig. 2F). Overall, the self-reports reflected an increase in anxiety (SA) and distress (SUDS) with the application of the stressors.

**Figure 2:**
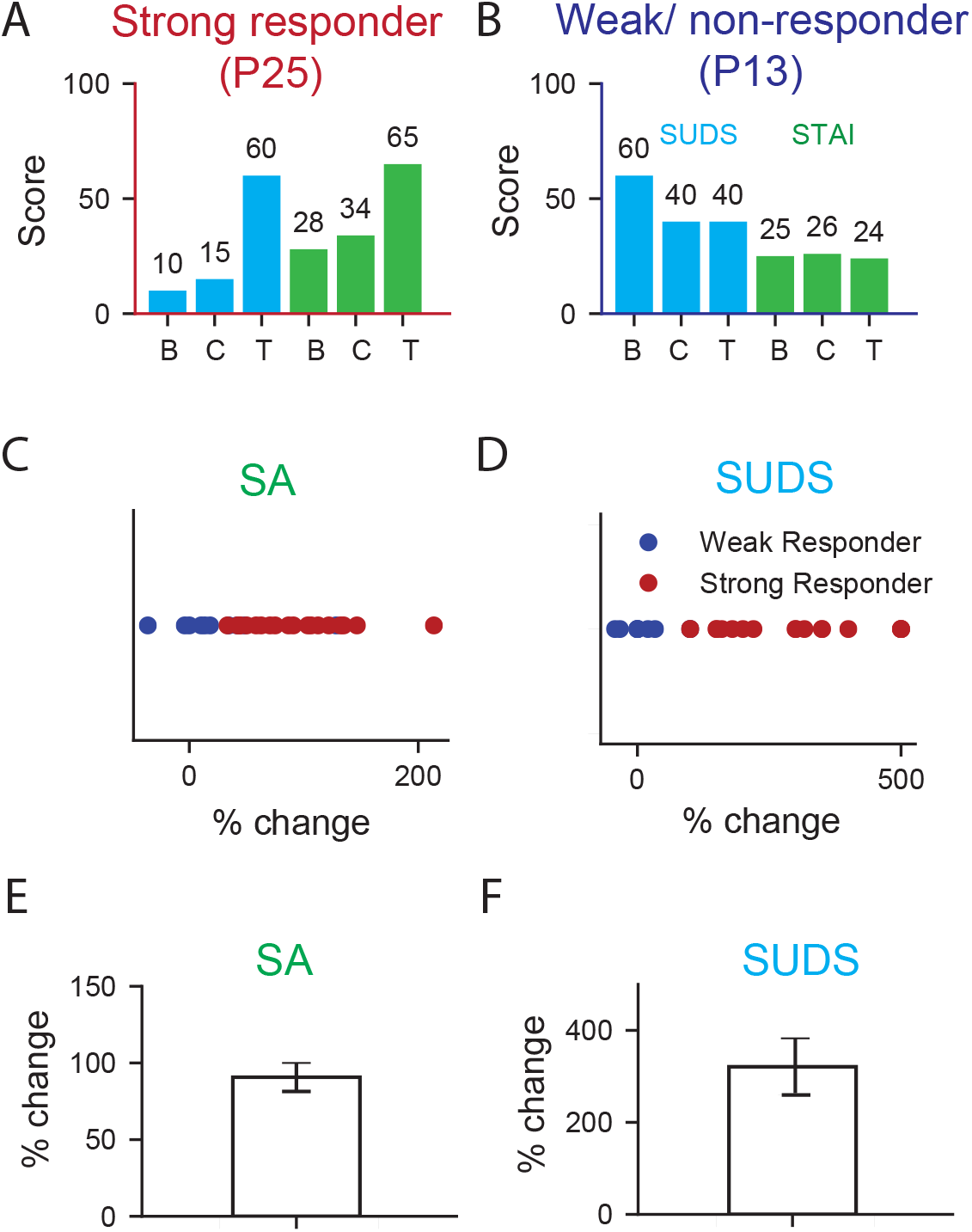
Strong and weak stress responders. A-B) Bar plots showing the SUDS scores (cyan) and state anxiety scores (SA, green) for the different epochs -- B- baseline, C- post-Cold pressor (CPT), T – post-Trier social stress test (TSST) – for an example strong responder in A (P25) and a weak/ non-responder in B (P13). Percentage change is state anxiety scores (C) and SUDS scores (in D) is shown for all strong responders (blue dots) and weak responders (red dots). An arbitrary threshold (STAI - 30%; SUDS – 50%) was chosen for illustrative purposes. The average change in SA post-TSST for the population (90±9%, N=23) is shown in (E). The average change in SUDS scores (320±61%) post-TSST is shown in (F).

### Physiological response to stressors

Electrodermal activity (EDA) refers to variation in the electrical properties of the skin in response to sweat secretion. By applying a low constant voltage, the change in skin conductance -- reflective of sympathetic tone -- can be measured non-invasively (45). Application of CPT and TSST induces autonomic arousal and increases sympathetic tone, which is expected to increase skin conductance (46). Change in EDA with respect to baseline (BC) is plotted in Fig 3A (top panel) for an example individual (P25, dashed red) and the group (solid black with confidence bands). As expected, P25 showed an elevation in EDA post-CPT compared to baseline (5 min post-CPT: 0.11 μS) that further increased post-TSST (5 min post-TSST: 0.13 μS). The group average also showed an increase in EDA with the application of the stressors (post CPT: 0.05 μS; post TSST: 0.09 μS). We then examined the increase in EDA post-CPT and post-TSST and compared that with the change in self-reported anxiety levels. EDA increased post-CPT with increase in anxiety, although the relationship was not statistically significant (r = 0.28, p=0.15, n=39). Post-TSST the relationship between EDA and anxiety remained nearly the same (Fig 3A, bottom panel, r = .26, p = .17, n = 39).

**Figure 3:**
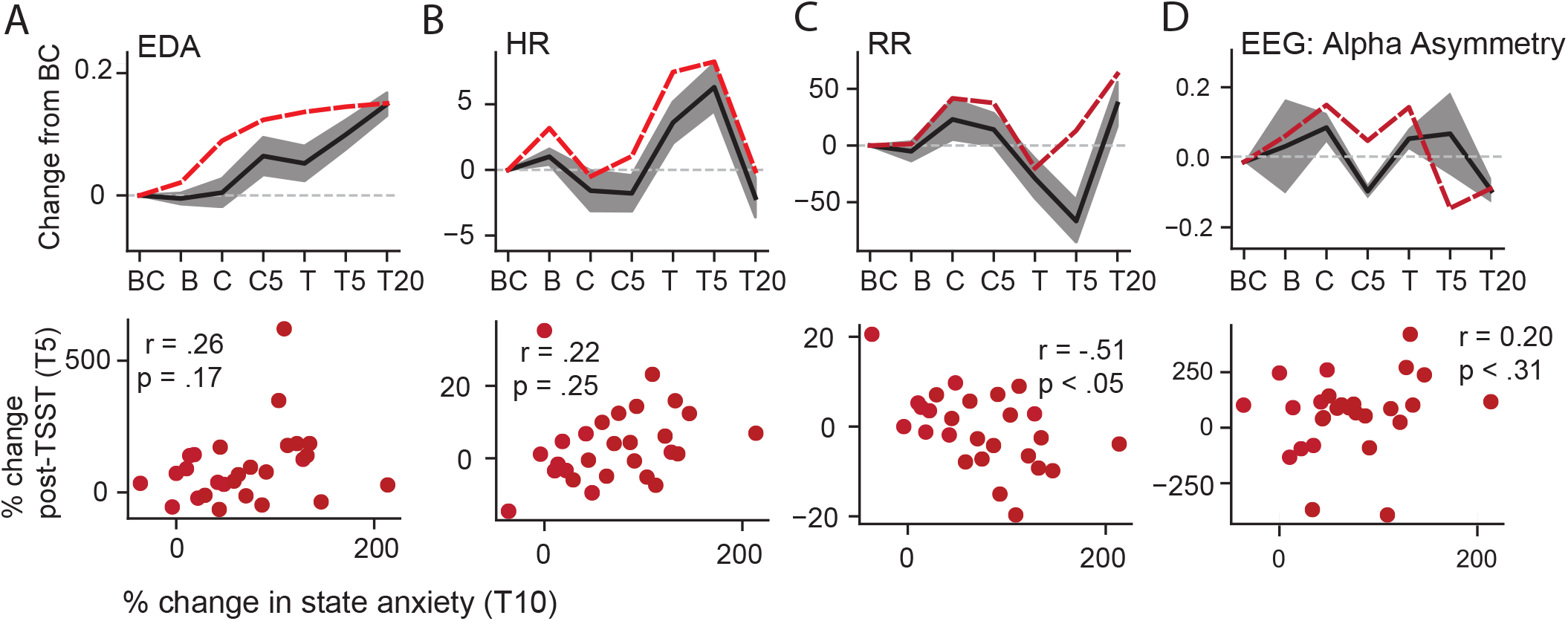
Physiological response to stressors. Top row of panels depict changes in electrodermal activity (EDA in μS, A), heart rate (HR in bpm, B), heart rate variability-based on R-R interval (RR – IBI in ms, C) and EEG alpha asymmetry (D). The x-axis shows the different epochs. BC- baseline control, B- baseline, C- onset of CPT, C5 – 5 min after C, T – onset of TSST, T5 and T20 – 5 and 20 minutes after T. Red dashed line is an example individual (P25). Black solid line and the confidence (standard error) indicates the group response. The y-axis reflects percentage change in the physiological response from BC: that is, change from BC divided by BC, times 100. In the bottom row of panels, the percentage change at T5 for individual participants is shown along with percentage change in state anxiety post-TSST onset (T10). Each red dot represents one participant. Pearson’s correlation and the associated p-values are indicated.

Acute stress increases sympathetic tone, which increases heart rate (HR) and decreases heart rate variability (HRV)(47). R-R intervals were extracted from a photoplethysmogram and HR and HRV were extracted (refer to Methods and Materials). Percentage change in HR (Fig 3B, top panel) and HRV (Fig 3C, top panel) with respect to a baseline (BC) is plotted for an example individual (P25, dashed red) and the group (solid black with confidence bands). P25 showed a slight decrease in HR post-CPT (5 min post-CPT: 1.04 bpm) and a strong increase post-TSST (5 min post-TSST: 8.24 bpm). At the same time, HRV post-CPT increased by 37.52 ms followed by a strong decrease post-TSST (13.14 ms, Fig 3C top panel). On average, the group showed a decrease in HR (−1.78 bpm) and an increase in HRV (14.10 ms) post-CPT. However, post-TSST the group on average displayed a robust increase in HR (6.3 bpm) accompanied by a decrease in HRV (66.61 ms). We then examined the change in HR and HRV post-CPT and compared that with the change in self-reported anxiety levels. Increase in HR (r = 0.27, p=0.16, n=39) and decrease in HRV (r = −0.35, p=0.06, n=39) were both correlated with change in state anxiety. We then plotted the change in HR (Fig 3B, bottom panel) and HRV (Fig 3C, bottom panel) post-TSST against the increase in self-reported change in state anxiety. The change in HR post-TSST was positively, although not significantly, correlated with change in self-reported stress (r = .22, p = .25, n = 39). Changes in HRV post-TSST relative to baseline were negatively correlated with changes in self-reported stress (r = −0.51, p < 0.05, n = 39). In summary, in our sample, HR increased and HRV decreased with increase in self-reported state anxiety following stressor application.

We finally examined EEG responses to acute stress. In particular, we plotted the difference in alpha activity between left and right frontal regions of the brain (frontal alpha asymmetry (FAS) (48)). Greater left than right frontal alpha (F3 > F4) characterizes approach-oriented situations whereas greater right than left frontal alpha (F4 > F3) is thought to reflect withdrawal-related motivational traits and states (49–51). Following CPT and TSST we expected a greater right frontal alpha asymmetry, i.e. F4 > F3. For the example individual (P25), FAS (log F4 – log F3) increased as expected both during CPT (0.16) and during TSST (0.16) compared to baseline (Fig. 3D, top panel). For the group, FAS increased on average post-CPT by 0.8 and post-TSST by 0.56 (Fig. 3D, top panel). We then compared the increase in FAS post-CPT and post-TSST with the change in self-reported anxiety levels. Increase in FAS Post-CPT was correlated with increase in anxiety but the trend was not significant (r = 0.14, p=0.49, n=39). Post-TSST the relationship between FAS and anxiety became stronger but was still not significant (Fig 3D, bottom panel, r = 0.20, p < 0.31, n = 39).

In summary, individual participant’s EDA, HR, HRV and FAS responded to the two stressors. Although changes in these measures reflected changes in self-reported state anxiety, this relationship did not reach statistical significance. This could partly be due to mismatch between physiology and self-report (See Discussion). To address this possibility, in the next section we describe how these measures can be combined using machine learning algorithms to decode stressful epochs, which can be defined objectively.

### Developing and testing a machine learning algorithm to predict stress state

We tested two qualitatively different stressor application protocols (CPT and TSST) and engaged participants in five different ‘control’ epochs -- baseline rest epoch, a rest epoch between stressors, a simple decision making game before and after stress induction, and a ‘talking’ epoch (refer to Methods section for more details). For this analysis, the algorithm was simply trained to distinguish stressor application epochs from control epochs. We refer to this algorithm as the “stress state classifier.”

First, we compared the accuracy with which the classifier could use features from a single physiological measure (EDA, HRV or EEG) to identify a stress application epoch from a control epoch. The features derived from these measures are listed in Supplemental Table 3. Figure 4A shows the prediction accuracy for EEG (purple band), HRV (brown band) and EDA (green band). The x-axis plots the proportion of data utilized for training the algorithm. For example, 80% indicates that out of all the epochs from all the subjects, a random 80% of the epochs were used to train the algorithm while the remaining 20% were used for testing. As expected, as the classifier was trained on more data (50%-80%) prediction accuracy increased, for all physiological measures (Fig. 4A and Supplemental Table 2: Sensitivity, specificity and accuracy). EEG-based measures yielded classifiers with the highest accuracy. For example, when 80% of the data was used for training the classifier using EEG, mean accuracy was 74.5% (Across subjects: min=72%, max = 79%, N=39). When HRV was used, mean classifier accuracy dropped to 70.7% (min=67%, max = 74%, N=39), while for EDA classifier accuracy dropped to 67.2% (min=66%, max = 68%, N=39).

**Figure 4:**
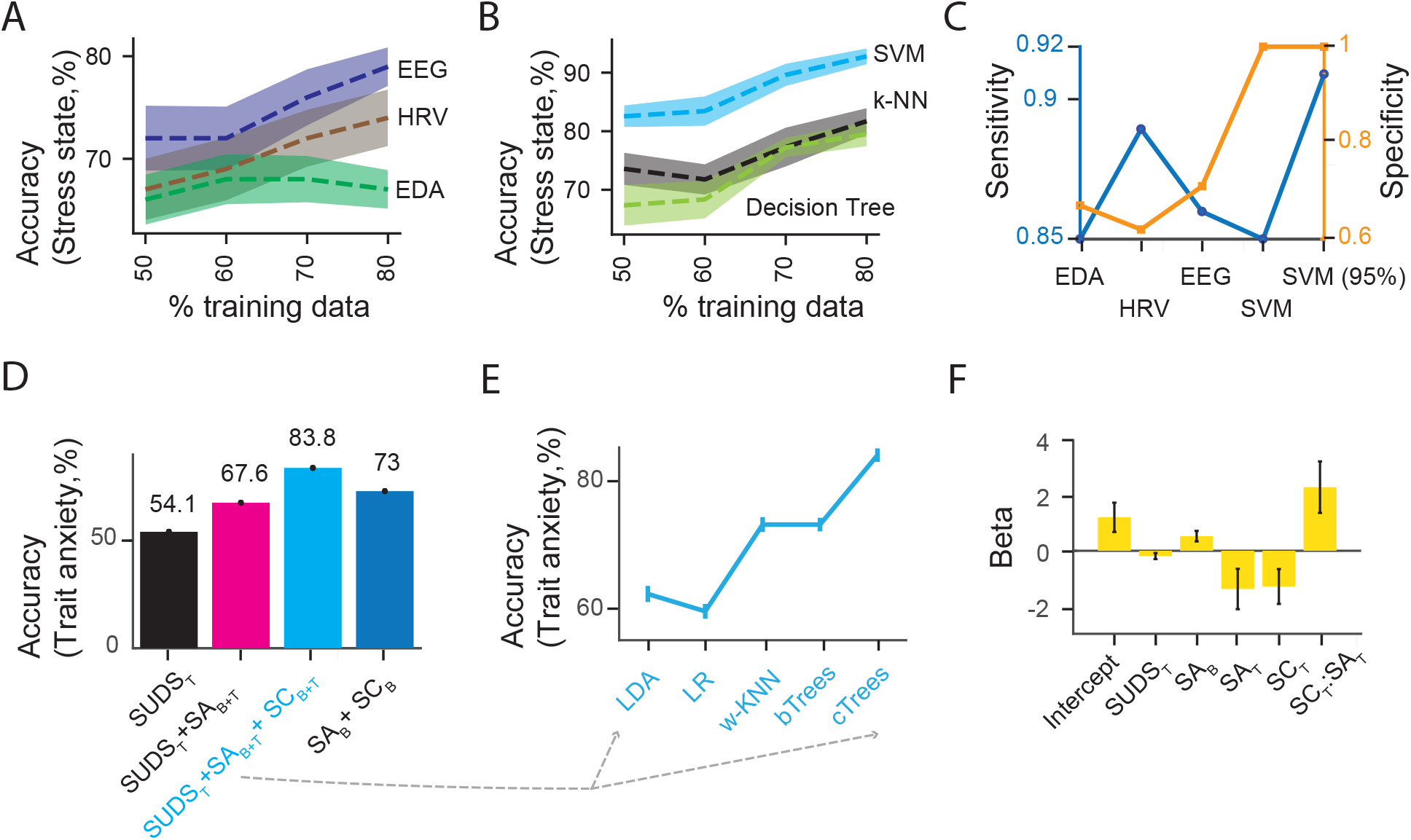
Identifying stress states and individuals with high trait anxiety. Prediction accuracy (A-B), sensitivity and specificity (C) of the stress state classification. Physiological measures -- EDA, HRV and EEG – were used independently in (A), whereas the best features from these measures were combined in (B), to identify stress states. Support vector machine (SVM), k-NN – k nearest neighbors and decision trees algorithms were used amongst others (Supplemental Table 4). The average prediction accuracy across participants is indicated by the bold dashed line, the standard error is indicated by the band around it. The percentage of the epochs utilized to train the algorithm – 50-80% -- is indicated on the x-axis. 80%of the data was used for training in (C) by default; SVM (95%) used 95% of the data for training. D) Average prediction accuracy across participants for those with high trait anxiety (> 35) using SUDS scores post-TSST (SUDS_T_), state anxiety scores at baseline (SA_B_) and post-TSST (SA_T_), stress state classifier scores at baseline (SC_B_) and post-TSST(SC_T_). These measures were combined optimally using complex trees (cTrees) in D. cTrees were compared with linear discriminant analysis (LDA), logistic regression (LR), weighted k-nearest neighbors (w-KNN) and boosted Trees (bTrees) in E. Coefficients (Beta±SE) obtained from a mixed generalized linear model are plotted to understand the relationship between SUDS_T_, SA_B_, SA_T_, SC_T_, the interaction terms, specifically, SA_T_ and SC_T_, and self-reported trait anxiety scores.

Next, best features were selected from the four independent streams of physiological data to train supervised machine learning algorithms (Supplemental Table 4: All algorithms). Figure 4B shows prediction accuracy for three of the algorithms tested. Performance of the SVM (blue band), KNN (black band) and decision trees (yellow band) was compared as a function of the proportion of data used for training vs testing (50-80%). Prediction accuracy increased with the size of the training set. SVM predicted stress epochs with the highest accuracy. For example, when 80% of the data was used for training, prediction accuracy of SVM was on average 92.2% (Fig 4B: Range: min = 81.9, max = 97.1, N=39), followed by decision trees at 81.3% (Range: min = 70.9, max = 90.9) and KNN at 77.8% (Range: min = 69.6, max = 90.9). Prediction accuracy of SVM further increased up to 95.8% when a larger fraction of the data was utilized to train the classifier (90% training data: SVM=94.5%, KNN=79.9%, Decision trees: 79.1%; 95% training data: SVM: 95.8%, KNN: 81.2%, Decision trees: 83.8%).

The sensitivity and specificity of the physiological measures and algorithms in identifying a stress state is plotted in Fig. 4C. Amongst the individual modalities, EEG-based predictions achieved highest specificity (EEG – 0.71, HRV – 0.62, EDA – 0.67) and HRV-based predictions achieved the highest sensitivity (EEG – 0.86, HRV – 0.89, EDA – 0.85). Combining the individual physiological measures using a SVM improved both sensitivity and specificity of the predictions. For example, when 95% of the data was utilized for training, SVM-based predictions achieved 91% sensitivity and 100% specificity (Fig. 4C).

Next, we tested the ability of the algorithm to predict the stress state of an individual participant whose data was fully withheld from the algorithm during training. To do this, we adopted the leave-one-out cross validation procedure. That is, one randomly-chosen participant’s data was left out and the algorithm was trained on the remaining 38 participants’ data. The trained algorithm was then used to forecast the stress states of the ‘left-out’ participant. This procedure was iteratively applied to all the participants. Stress state prediction accuracy declined but was still significantly above chance (SVM: Range: min = 77.7, max = 80.0, mean = 79.92, KNN: Range: min = 50.0, max = 80, mean = 71.2 and decisions trees: Range: min = 30.0, max = 90.0, mean = 66.8).

These findings indicate that amongst the physiological measures, EEG-based metrics predicted stressful epochs with higher accuracy than other physiological measures. Combining features from different physiological measures -- EEG, HRV, HR and EDA – improved stress state prediction accuracy relative to any single physiological channel. Our stress state classifier decoded stressful epochs significantly above chance in both out-of-sample epochs and individuals. Lastly, self-reported increases in anxiety post-TSST were consistent with the predictions of the stress state classifier for most participants (26/39). A fraction of the participants did not show alignment of self-report and physiological responses to stress induction. For 8/39 participants, the stress state classifier predicted an increase in stress, but self-reports did not. Conversely, 5/39 participants reported an increase in stress but the classifier estimate did not.

### Identifying individuals with high trait anxiety

Next, we tested whether self-report and physiological responses to stress manipulation could be used to identify individuals with high trait anxiety. To do this, we first median-split the sample into high trait anxiety (STAI-T > 35) and low trait anxiety (STAI-T<35). Then we trained a classifier (complex tree in Fig. 4D; other classifiers in Fig. 4E) using a leave-one-out cross validation procedure (other combinations of training set are tabulated in Supplementary Table 5).

That is, the algorithm was trained on all but one participant’s SUDS scores, state anxiety scores (SA), and the stress state classifier (SC) outputs assessed both at baseline and after TSST. Prediction accuracy using baseline SUDS scores (51.4%; Sensitivity = 72%; Specificity = 32%) was near chance levels (50%). Including SUDS post-TSST increased prediction accuracy to 54.1% (Sensitivity = 28%; Specificity = 79%). Adding baseline and post-TSST STAI-S reports improved prediction accuracy to 67.6% (Sensitivity = 72%; Specificity = 63%). Including the estimates from the stress state classifier further improved prediction accuracy to 83.8% (Sensitivity = 94%; Specificity = 74%). When only baseline SA and stress state classifier outputs were used, prediction accuracy dropped slightly but was still well above chance at 73% (Sensitivity = 78%; Specificity = 68%). These results show that stress state classifier estimates based on physiological measures can improve the prediction accuracy of a high trait anxiety individual from 67.6 to 83.8 %, a 16.2% improvement. For this analysis, the complex trees classifier was the best-performing algorithm (Fig. 4E; complex trees = 83.8%; boosted trees =73%, weighted k-NN =73%, logistic regression = 59.5% and LDA =62.2%).

Next, to understand the relationship between SUDS, SA, stress state classifier (SC) estimates and trait anxiety, we developed a mixed generalized linear model (Fig. 4F). Trait anxiety scores served as the dependent variable, while SUDS, SA and stress classifier estimates -- both at baseline and post TSST – served as the independent variables (See Methods for details). Baseline SA (0.53±0.19, tstat=2.7, DF=29, p<0.009) was a significant predictor of trait anxiety. Post-TSST SA increase was also nearly significant as a predictor (−1.38±0.73, tstat= - 1.88, DF=29, p<0.068). Post-TSST stress state classifier estimates (−1.29±0.64, tstat=-2.02, DF=29, p<0.053) was also nearly significant; its interaction with post-TSST SA (2.31±0.93, tstat=2.4, DF=29, p<0.02) was a significant predictor of trait anxiety. Baseline and post-TSST SUDS (Baseline: 0.002±0.001, tstat=1.5, DF=26, p<0.12; post-TSST: −0.19±0.12, tstat= −1.58, DF=27, p<0.12), as well as baseline stress state classifier estimates (−0.23±1.02, tstat=-0.23, DF=27, p<0.81), were not significant predictors.

Results from the mixed model indicate that individuals with higher trait anxiety displayed increased state anxiety (SA) at baseline. Further, individuals with high trait anxiety displayed a muted response to acute stressors, as indicated by the negative coefficients for SA and stress state classifier, compared with individuals with low trait anxiety. The strong interaction effects between these predictors suggests they at least partially index the same underlying states. Notably, SUDS estimates were only weakly informative about trait anxiety. In summary, individuals with high trait anxiety can be identified with a machine learning approach that utilizes multiple physiological measures in conjunction with self-reports.

## Discussion

We examined the stress responses of healthy participants using self-reports and multiple physiological measures. Participants were exposed to 2 kinds of acute stressors and engaged in five non-stressful activities. During this experience, we observed robust changes in EEG, HRV, HR, GSR that were weakly correlated with increases in self-reported stress. By combining select features from these different measures using a machine learning-based algorithm, stressful states could be identified from non-stressful states with ~96% accuracy. Further, individuals with high trait anxiety were identified with ~84% accuracy using a decision tree to optimally combine physiological measures with self-reported state anxiety. While individuals with high trait anxiety displayed high baseline state anxiety, their response to acute stressors was muted.

### Identifying stressful states and non-stressful states

In this study, we used several physiological measures to understand individual responses to a stressor. GSR has been extensively used to measure stress response (22–29) and, even in this study, GSR increased with stress induction. However, GSR as a measure was not as sensitive as heart rate or EEG-based measurements in identifying stress states. HRV reliably declines with increases in stress (52–54) and in our study HRV also decreased with self-reported increases in stress-induced state anxiety. Nevertheless, HRV-based measures lacked the specificity of EEG. That is, many non-stressful epochs were wrongly identified as stressful epochs, limiting the utility of HRV in this context (55,56). Several recent studies have attempted to combine different physiological measures to improve the ability of an algorithm to identify stress states (24,25,29,33,38–42). By combining select features from these physiological modalities using a support vector machine, we achieved high sensitivity (91%) and specificity (100%)—indeed more than that achieve using any independent measure alone (Supplementary Table 2).

Two other factors contributed to the improvement in the prediction accuracy of the classifier. First, by including two different stressors and five different non-stressful epochs, the classifier was trained across a variety of different conditions varying in intensity, engagement, and arousal. Second, increasing the training set to >35 individuals enabled the classifier to remain robust to individual differences in stress response. Overall, by combining features from physiological measures to develop the classifier, and by training the classifier on a variety of different conditions and individuals, the classifier’s ability to accurately identify a stressful state in untested conditions and individuals improved. In light of these findings, we believe that a larger training set with more diverse conditions would not only help validate the current results but also improve the generalizability of the classifier.

In a majority of participants (26/39), self-reported responses to a stressor were consistent with physiology-based assessments. In other participants, however, there were discrepancies between self-report and physiology. Some participants did not report a response to the stressor but their physiology indicated otherwise. Such mismatches suggest that participants may be unaware of, or unable to access, or simply unwilling to report physiological responses to stress and other such internal states (57–59). By contrast, some participants reported a response to the stressor but their physiological measurements did not register it. These findings highlight the potential for inaccuracy or bias in self-report and the need for a more objective “ground truth” rooted in observable dysfunctions in daily life activities. Digital approaches to mental health, such as search terms in web browsing (60–62), financial behaviour (63), or other observable “big data” may provide a complementary source of information to refine our biometric approach.

### Recognizing individuals with high trait anxiety

Here we studied a nonclinical population by obtaining self-reported anxiety characteristics using measures like the State-Trait Anxiety Inventory (11), which we used to identify individuals with high trait anxiety. Critical differences exist between high trait anxiety and clinical anxiety disorders, but the frequency of conversion from elevated trait anxiety to clinical disorders suggests our findings could provide important insights into anxiety disorders (64). Unmanaged or chronic stress has been implicated in the process of conversion to anxiety disorders. Understanding the relationship between early-life stressful events or chronic, unmanaged stress may help uncover the mechanisms underlying the onset of anxiety disorders (65).

Different personality types may vary considerably in how they respond to stressful events. Evolutionary biologists have reasoned that such differences are maintained in the population because they support frequency-dependent adaptation under different conditions. Hawks and Doves are two phenotypes that have been related to variations in HPA axis activity and anxiety (66). In response to threat, Doves, who are typically anxious, shy and cautious, adopt a flight strategy. They also maintain a high baseline HPA axis response due to increased anxiety, as well as low sympathetic and adrenomedullary response to acute threats. Hawks, on the other hand, display highly aggressive behavior and preferentially adopt a fight strategy. They exhibit low baseline HPA axis response but display a high sympathetic and adrenomedullary responses to acute threats(66–68). In our study, a blunted response to acute stressors in high trait anxiety participants is consistent with a “dove-like” phenotype (68,69). From a more mechanistic perspective, these observations suggest that by manipulating stress and by measuring selfreported changes in state anxiety levels and the associated changes in physiology, one can determine biomarkers of specific phenotypes.

Overall, our study provides an approach combining self-reports with physiological responses to stress to identify individuals with high trait anxiety. Accurate and continuous stress detection is a valuable facet of any treatment plan for a broad range of anxiety and stress-related disorders. Precisely measuring incremental progress through stressful experiences can boost patient’s knowledge, confidence, and self-efficacy, which studies have shown improves mental health treatment outcomes. Specifically, improvements in self-efficacy are associated with improvements in confidence to maintain lifestyle changes, which is a key component of stress management (70). Overall, these results demonstrate the feasibility of using machine learning analyses of biometric data to enable sensitive diagnosis and continuous monitoring of mental health conditions like anxiety and depression.

## Methods

### Participants

Forty (40) participants were recruited between the ages of 18 and 35 (mean = 25.15 / SD = 5.04; 43.5% Female / 56.4% Male) with no diagnosed psychiatric disorders or psychotropic medication use (based on self-report). Participants were recruited from the local community through the Wharton Behavioral Lab. Basic non-PHI demographic information (e.g., age, sex, level of education) was collected and a unique subject ID number was assigned to each participant (Supplementary Table 1). Participants were instructed to not engage in exercise or consume caffeinated beverages for a minimum of two hours prior to their scheduled visit time. Study personnel explained the purpose, potential risks of the experiment and completed the informed consent process with each participant following protocols approved by the University of Pennsylvania’s Institutional Review Board (IRB) in compliance with the Helsinki Declaration. All participants gave written informed consent filed with the University of Pennsylvania’s IRB.

At baseline, participants were outfitted with a wireless EEG headset (microEEG, Bio-Signal Group Corp., Acton, MA) and multiple HR/GSR sensors (Shimmer GSR+, Shimmer Inc., Boston, MA; E4 Wristband, Empatica Inc., Boston, MA). iMotions (Copenhagen, Denmark) was used to synchronize all the devices.

### Procedure

Trait anxiety survey was administered online prior to the visit via Qualtrics. On the session day, set-up, and calibration was carried out upon arrival. Following initial briefing and set-up, participants were asked to relax for 10 minutes. The first 5 minutes were considered baseline control (BC) and the physiological readings during the session were measured relative to this epoch. The next 5 minutes were considered baseline period (B). Participants’ self-reports - both subjective units of distress (SUDS) and state anxiety (SA) -- were obtained during the baseline period. Then, participants played a simple video game for 10 minutes after 2 minutes of practice. In the game, they picked berries from a virtual berry patch (71–74). Then the cold pressor test (CPT) was administered to induce temporary acute physical stress. During this task participants were provided with an arm wrap made of cold gel packs (35-40 °F)(75). The cold gel packs were placed around their dominant forearm for three minutes. The wrap is designed to be stressful and uncomfortable but not painful or dangerous(76,77). Following this task, participants were instructed to quietly rest for 6 minutes. Second set of self-reports – both SUDS and SA -- were obtained at the beginning of the rest period. After the quiet rest, participants were asked to talk about something or engaged in casual conversation for three minutes. Following this, the Trier Social Stress Test (TSST) was administered(78). In this epoch the participants were asked to prepare and deliver a short speech to a socially evaluative audience(79–81). Participants were provided with pen and paper to take notes and informed that they had three minutes to prepare a five-minute speech about interviewing for their ideal job (step 1). To increase stress, they were instructed that their responses would be recorded (audio only if they did not consent to video) and that an evaluator trained in public speaking would judge their speech for honesty, believability, and confidence. Prior to giving their speech, the participant’s notes were taken away and a five minute timer was started (step 2). If they stopped talking before the five minutes ended, they were told by the research coordinator: “You still have time remaining, please keep talking about why you should be hired for this job.” At the end of the speech, the evaluator provided the participants with the following instructions (step 3): “During the final one minute you will be asked to sequentially subtract the number 13 from 1,022. You will verbally report your answers aloud, and be asked to start over from 1,022 if a mistake is made. Your time begins now.” If the participant made an error, they were told “That is incorrect, please start over from 1,022.” Third set of self-reports – both SUDS and SA -- were obtained soon after TSST. After that participants played the patch-foraging video game again for 10 minutes. After this, they were debriefed and the equipment was removed.

Participants completed the standardized Spielberger’s state anxiety Form Y as a report for state anxiety (SA) and rated their distress level -- Subjective Units of Distress (SUDs) -- on a 0-100 scale (0 = calm and free from distress and 100 = the most distressed they can ever recall being) (82).

One participant quit the study midway after the CPT. This participant’s data was excluded from data analysis.

### Data Analysis

All data processing and statistical analyses were carried out on MATLAB and Python.

State-Trait Anxiety Inventory (STAI): All participants completed the STAI (11), which was used for assessing both trait and state anxiety. There are 20 items for assessing trait anxiety and 20 for state anxiety. All items are rated on a 4-point scale, with higher scores indicating greater anxiety.

Subjective Units of Distress Scale (SUDS): Participants completed the SUDS both at baseline and following each stress induction task. The SUDS is a widely used tool for assessing the subjective intensity of distress and other internal experiences, such as anxiety, anger, and agitation. The scale ranges from 1 to 100, with 100 signifying the most intense feelings.

HR/Heart Rate Variability (HR/HRV): Heart rate data was obtained throughout the duration of each session through the use of an E4 Wristband. The E4 is equipped with a PPG sensor which measures blood volume pulse (BVP). From this measurement both HR and HRV were derived. HRV is the beat-to-beat alteration in heart rate that can be used as a noninvasive biomarker for autonomic nervous system activity (83). Standard time and frequency domain derivatives of HRV were calculated.

Electrodermal Activity (EDA) / Galvanic Skin Response (GSR): Electrodermal activity (EDA) was continuously recorded throughout the duration of each session through the use of a Shimmer GSR+. The GSR sensor monitors skin conductivity which reflects the variations in the electrical characteristics of the skin. Skin conductance is modulated by sympathetic activity and it is directly correlated with emotional arousal.

EEG Activity: microEEG was used to obtain EEG signals throughout the duration of the session (84). Signals were collected at a 250 Hz sampling rate from electrode channels along the scalp. EEG signals were first re-referenced to the average of the two earlobes and then filtered between 1 and 50 Hz. Next, epochs were extracted and probabilistically improbable data points (3 SD from mean) were detected as artifacts and removed. Additional artifact detection to remove eye blink and movement artifact components was subsequently performed using independent component analysis. All the preprocessing was done using EEGLAB (85).

For each epoch, Welch’s power spectral density estimate was used to transform data from the time domain to the frequency domain in order to decompose the signal and calculate absolute and relative band powers, asymmetries, and coherence in addition to the phase of the EEG channel signals. Band powers calculated included delta (1-4 Hz), theta (4-8 Hz), alpha (8-13 Hz), low beta (13-25 Hz), high beta (25-30 Hz), and gamma (30-50 Hz) from the prefrontal, frontal, temporal, parietal, and occipital brain regions.

### Classifier design

Following preprocessing, different features were extracted from the HR and EEG data (Supplement table 3). Average data (1 to 3 min) from different epochs (stress vs control) were then used to train the classifier. Different machine learning algorithms (Supplementary table 2) were trained on the data streams. Data was upsampled to ensure class balance. Dataset was split up into partitions (2–10), where data from all-but-one partitions were used as the training set and left-out data was used as the testing set. This was repeated for all partitions, so that each epoch was part of the test set at least once. Cross-validation also helped to detect and limit overfitting. Feature selection was carried out to reduce the feature set size. Grid search was implemented to optimize the parameters used. Grid search and feature selection were carried out on the training dataset prior to testing. To diagnose bias and variance in the models’ outcome, area under the ROC curves were evaluated. The primary metrics used to indicate each model’s performance were accuracy, sensitivity, and specificity.

For the classifier implemented to identify individuals with trait anxiety, leave-one-out cross-validation approach was followed. In this case one participant’s data was left out of the training dataset. The remaining 38 participants’ data was used to train the different algorithms. Other training and testing set ratios (90-10, 95-5) were also tested (Supplementary Table 5).

### Generalized linear model with mixed effects

To take advantage of the continuous nature of the trait anxiety scores, we developed generalized linear model with mixed effects.

The following model was tested:

Trait anxiety ~ SUDS_B_ + SUDS_T_ + STAI_B_ + STAI_T_ + SC_B_ + SC_T_ + STAI_T_: SC_T_ + (1 | participant)

SUDS_B_ - Baseline SUDS, SUDS_T_ - post-TSST SUDS, SA_B_ - Baseline state anxiety, SA_T_ - state anxiety post-TSST, SC_B_ - Baseline stress Classifier output, SC_T_ – post-TSST stress Classifier output are the fixed effects terms. “1| participant” is the random intercept that accounted for differences between participants.

Other combinations – including/excluding fixed effect terms, including other interactions terms for fixed effects, including random slopes for the participants, etc, were also tested.

Since the trait anxiety score distribution is right skewed and all-positive, it was modelled using a “Gamma” distribution with an “identity” link function. All analyses were carried out in Matlab using the “fitglme” function.

## Acknowledgements

We thank the Wharton Behavioral Lab and Robert Botto for help with setting up the experiment; Magdalena Araya Curutchet, Ingrid Tous, Leeka Ran, Jici Wang, Felipe Parodi for support with data collection; James Henson and Samah Baki of Biosignal group Inc for providing us with the EEG device and excellent technical support; and Javier Omar and Dhaval Bhatt for help with manuscript preparation.

The study was partially funded by NeuroFlow and the Wharton Neuroscience Initiative. AR was supported in part by the P&S fund and the Brain and Behavior Research Foundation. MLP was supported in part by NIH grants R37-MH-109728-01, R01-MH-108627-01 and the Simon’s Foundation.

## Author contributions

AR, AP, CM and MLP conceived the project. AR and AP designed the experiment and supervised data collection. AR and WL performed data analysis. AR and MLP wrote the manuscript.

## Supplementary Tables

Table 1: Demographics

Table 2: Accuracy, Sensitivity and Specificity of the physiological measures and algorithms

Table 3: Features derived from physiological measurements

Table 4: Stress state classifier: List of all machine learning algorithms trained and tested

Table 5: Identifying individuals with high trait anxiety. Different training sets and algorithms tested.

## References

1. Managing Stress and Anxiety | Anxiety and Depression Association of America, ADAA.

2. Kessler RC. Posttraumatic stress disorder: The burden to the individual and to society. J Clin Psychiatry. 2000;

3. Kessler RC, Berglund P, Demler O, Jin R, Merikangas KR, Walters EE. Lifetime prevalence and age-of-onset distributions of DSM-IV disorders in the national comorbidity survey replication. Arch Gen Psychiatry. 2005;

4. Bandelow B, Michaelis S. Epidemiology of anxiety disorders in the 21st century. Dialogues Clin Neurosci. 2015;

5. Evans TM, Bira L, Gastelum JB, Weiss LT, Vanderford NL. Evidence for a mental health crisis in graduate education. Nat Biotechnol. 2018;

6. John-Baptiste AA, Li L, Isaranuwatchai W, Osuch E, Anderson KK. Healthcare utilization costs of emerging adults with mood and anxiety disorders in an early intervention treatment program compared to a matched cohort. Early Intervention in Psychiatry. 2019;

7. Grupe DW. Decision-making in anxiety and its disorders. Decis Neurosci An Integr Perspect. 2016;327–38.

8. American Psychiatric Association. Diagnostic and Statistical Manual of Mental Disorders (5th Edition). American Journal of Psychiatry. 2013.

9. Raymond JG, Steele JD, Seriès P. Modeling trait anxiety: From computational processes to personality. Frontiers in Psychiatry. 2017.

10. Shackman AJ, Tromp DPM, Stockbridge MD, Kaplan CM, Tillman RM, Fox AS. Dispositional negativity: An integrative psychological and neurobiological perspective. Psychol Bull. 2016;

11. Spielberger CD. State-trait anxiety inventory: a comprehensive bibliography. Consult Psychol Press. 1989;

12. McEwen BS. Physiology and neurobiology of stress and adaptation: central role of the brain. Physiol Rev. 2007;

13. Robins LN. Epidemiology: Reflections on Testing the Validity of Psychiatric Interviews. Arch Gen Psychiatry. 1985;

14. Knowles ES, Coker MC, Scott RA, Cook DA, Neville JW. Measurement-Induced Improvement in Anxiety: Mean Shifts with Repeated Assessment. J Pers Soc Psychol. 1996;

15. Sharpe JP, Gilbert DG. Effects of repeated administration of the beck depression inventory and other measures of negative mood states. Pers Individ Dif. 1998;

16. Windle C. Test-Retest Effect on Personality Questionnaires. Educ Psychol Meas. 2007;

17. Shrout PE, Stadler G, Lane SP, McClure MJ, Jackson GL, Clavél FD, et al. Initial elevation bias in subjective reports. Proc Natl Acad Sci. 2018;

18. Allen AP, Kennedy PJ, Cryan JF, Dinan TG, Clarke G. Biological and psychological markers of stress in humans: Focus on the Trier Social Stress Test. Neuroscience and Biobehavioral Reviews. 2014.

19. Campbell J, Ehlert U. Acute psychosocial stress: Does the emotional stress response correspond with physiological responses? Psychoneuroendocrinology. 2012.

20. Nater UM, Ditzen B, Strahler J, Ehlert U. Effects of orthostasis on endocrine responses to psychosocial stress. Int J Psychophysiol. 2013;

21. Urwyler SA, Schuetz P, Sailer C, Christ-Crain M. Copeptin as a stress marker prior and after a written examination-the CoEXAM study. Stress. 2015;

22. Healey JA, Picard RW. Detecting stress during real-world driving tasks using physiological sensors. IEEE Trans Intell Transp Syst. 2005;

23. Zhai J, Barreto A. Stress detection in computer users through non-invasive monitoring of physiological signals. In: Biomedical Sciences Instrumentation. 2006.

24. De Santos Sierra A, Sánchez Ávila C, Guerra Casanova J, Bailador Del Pozo G. A stress-detection system based on physiological signals and fuzzy logic. IEEE Trans Ind Electron. 2011;

25. Katsis CD, Katertsidis NS, Fotiadis DI. An integrated system based on physiological signals for the assessment of affective states in patients with anxiety disorders. In: Biomedical Signal Processing and Control. 2011.

26. Willmann M, Langlet C, Hainaut JP, Bolmont B. The time course of autonomic parameters and muscle tension during recovery following a moderate cognitive stressor: Dependency on trait anxiety level. Int J Psychophysiol. 2012;

27. Karthikeyan P, Murugappan M, Yaacob S. Multiple physiological signal-based human stress identification using non-linear classifiers. Elektron ir Elektrotechnika. 2013;

28. Seoane F, Mohino-Herranz I, Ferreira J, Alvarez L, Buendia R, Ayllón D, et al. Wearable biomedical measurement systems for assessment of mental stress of combatants in real time. Sensors (Switzerland). 2014;

29. Kukolja D, Popović S, Horvat M, Kovač B, Ćosić K. Comparative analysis of emotion estimation methods based on physiological measurements for real-time applications. Int J Hum Comput Stud. 2014;

30. Wijsman J, Grundlehner B, Liu H, Penders J, Hermens H. Wearable physiological sensors reflect mental stress state in office-like situations. In: Proceedings - 2013 Humaine Association Conference on Affective Computing and Intelligent Interaction, ACII 2013. 2013.

31. Thayer JF, Åhs F, Fredrikson M, Sollers JJ, Wager TD. A metaanalysis of heart rate variability and neuroimaging studies: Implications for heart rate variability as a marker of stress and health. Neuroscience and Biobehavioral Reviews. 2012.

32. Soares-Caldeira LF, De Souza EA, De Freitas VH, De Moraes SMF, Leicht AS, Nakamura FY. Effects of additional repeated sprint training during preseason on performance, heart rate variability, and stress symptoms in futsal players: A randomized controlled trial. J Strength Cond Res. 2014;

33. Al-Shargie F, Kiguchi M, Badruddin N, Dass SC, Hani AFM, Tang TB. Mental stress assessment using simultaneous measurement of EEG and fNIRS. Biomed Opt Express. 2016;

34. Mühl C, Jeunet C, Lotte F. EEG-based workload estimation across affective contexts. Front Neurosci. 2014;

35. Alonso JF, Romero S, Ballester MR, Antonijoan RM, Mañanas MA. Stress assessment based on EEG univariate features and functional connectivity measures. Physiol Meas. 2015;

36. Giles GE, Mahoney CR, Brunyé TT, Taylor HA, Kanarek RB. Stress effects on mood, HPA axis, and autonomic response: Comparison of three psychosocial stress paradigms. PLoS One. 2014;

37. Valdés A. Measurement of acute psychological stress. 2017.

38. Sharma N, Gedeon T. Hybrid genetic algorithms for stress recognition in reading. In: Lecture Notes in Computer Science (including subseries Lecture Notes in Artificial Intelligence and Lecture Notes in Bioinformatics). 2013.

39. Kurniawan H, Maslov A V, Pechenizkiy M. Stress detection from speech and galvanic skin response signals. In: Proceedings of CBMS 2013 - 26th IEEE International Symposium on Computer-Based Medical Systems. 2013.

40. Van Den Broek EL, Van Der Sluis F, Dijkstra T. Cross-validation of bimodal health-related stress assessment. Pers Ubiquitous Comput. 2013;

41. Ghaderi A, Frounchi J, Farnam A. Machine learning-based signal processing using physiological signals for stress detection. 2015 22nd Iran Conf Biomed Eng ICBME 2015. 2016;

42. Keshan N, Parimi P V, Bichindaritz I. Machine learning for stress detection from ECG signals in automobile drivers. Proc - 2015 IEEE Int Conf Big Data, IEEE Big Data 2015. 2015;

43. Lovallo W. The Cold Pressor Test and Autonomic Function: A Review and Integration. Psychophysiology. 1975;

44. Mcrae AL, Saladin ME, Brady KT, Upadhyaya H, Back SE, Timmerman MA. Stress reactivity□: biological and subjective responses to the cold pressor and Trier Social stressors y. 2006;(January):377–85.

45. Fowles DC, Christie MJ, Edelberg R, Grings WW, Lykken DT, Venables PH. Publication Recommendations for Electrodermal Measurements. Psychophysiology. 1981;

46. Benedek M, Kaernbach C. A continuous measure of phasic electrodermal activity. J Neurosci Methods. 2010;

47. Mestanik M, Mestanikova A, Visnovcova Z, Calkovska A, Tonhajzerova I. Cardiovascular sympathetic arousal in response to different mental stressors. Physiol Res. 2015;

48. Davidson R, Schwartz C, Saron E, Bennett J, Goleman D. Frontal versus parietal EEG asymmetry during positive and negative affect. Psychophysiology. 1979;

49. Mathersul D, Williams LM, Hopkinson PJ, Kemp AH. Investigating Models of Affect: Relationships Among EEG Alpha Asymmetry, Depression, and Anxiety. Emotion. 2008;

50. De Pascalis V, Cozzuto G, Caprara GV, Alessandri G. Relations among EEG-alpha asymmetry, BIS/BAS, and dispositional optimism. Biol Psychol. 2013;

51. Harmon-Jones E, Gable PA, Peterson CK. The role of asymmetric frontal cortical activity in emotion-related phenomena: A review and update. Biological Psychology. 2010.

52. Dishman RK, Nakamura Y, Garcia ME, Thompson RW, Dunn AL, Blair SN. Heart rate variability, trait anxiety, and perceived stress among physically fit men and women. Int J Psychophysiol. 2000;

53. Acharya UR, Joseph KP, Kannathal N, Lim CM, Suri JS. Heart rate variability: A review. Med Biol Eng Comput. 2006;

54. Kim HG, Cheon EJ, Bai DS, Lee YH, Koo BH. Stress and heart rate variability: A meta-analysis and review of the literature. Psychiatry Investig. 2018;

55. Boonnithi S, Phongsuphap S. Comparison of heart rate variability measures for mental stress detection. 2011 Comput Cardiol. 2011;

56. Mackersie CL, Calderon-Moultrie N. Autonomic Nervous System Reactivity During Speech Repetition Tasks. Ear Hear. 2016;

57. Hunt M, Auriemma J, Cashaw ACA. Self-report bias and underreporting of depression on the BDI-II. J Pers Assess. 2003;

58. Stöber J. Reliability and validity of two widely-used worry questionnaires: Self-report and self-peer convergence. Pers Individ Dif. 1998;

59. Semmer NK, Grebner S, Elfering A. BEYOND SELF-REPORT: USING OBSERVATIONAL, PHYSIOLOGICAL, AND SITUATIONBASED MEASURES IN RESEARCH ON OCCUPATIONAL STRESS. Research in Occupational Stress and Well Being. 2003.

60. Wang Y, Kung LA, Byrd TA. Big data analytics: Understanding its capabilities and potential benefits for healthcare organizations. Technol Forecast Soc Change. 2018;

61. Raghupathi W, Raghupathi V. Big data analytics in healthcare: promise and potential. Heal Inf Sci Syst. 2014;

62. Powles J, Hodson H. Google DeepMind and healthcare in an age of algorithms. Health Technol (Berl). 2017;

63. www.Whealthcare.org.

64. Kendler KS, Gardner CO. A longitudinal etiologic model for symptoms of anxiety and depression in women. Psychol Med. 2011;

65. Raymond JG, Steele JD, Seriès P. Modeling trait anxiety: From computational processes to personality. Front Psychiatry. 2017;8(JAN):1–19.

66. Korte SM, Koolhaas JM, Wingfield JC, McEwen BS. The Darwinian concept of stress: benefits of allostasis and costs of allostatic load and the trade-offs in health and disease. Neurosci Biobehav Rev. 2005;

67. Ellis BJ, Jackson JJ, Boyce WT. The stress response systems: Universality and adaptive individual differences. Dev Rev. 2006;

68. Souza GGL, Mendonça-de-Souza ACF, Duarte AFA, Fischer NL, Souza WF, Coutinho E, et al. Blunted cardiac reactivity to psychological stress associated with higher trait anxiety: A study in peacekeepers. BMC Neurosci. 2015;

69. Peng H, Wu J, Sun X, Guan Q, Luo Y. Trait anxiety predicts the response to acute psychological stress. Acta Psychol Sin. 2018;

70. Ludman EJ, Peterson D, Katon WJ, Lin EHB, Von Korff M, Ciechanowski P, et al. Improving confidence for self care in patients with depression and chronic illnesses. Behav Med. 2013;

71. Addicott MA, Pearson JM, Kaiser N, Platt ML, Joseph McClernon F. Suboptimal foraging behavior: A new perspective on gambling. Behav Neurosci. 2015;

72. Addicott MA, Pearson JM, Sweitzer MM, Barack DL, Platt ML. A Primer on Foraging and the Explore/Exploit Trade-Off for Psychiatry Research. Neuropsychopharmacology [Internet]. 2017;(November 2016):1–33. Available from: http://www.nature.com/doifinder/10.1038/npp.2017.108

73. Hayden BY, Pearson JM, Platt ML. Neuronal basis of sequential foraging decisions in a patchy environment. Nat Neurosci [Internet]. 2011;14(7):933–9. Available from: http://dx.doi.org/10.1038/nn.2856

74. Ramakrishnan A, Hayden BY, Platt ML. Local field potentials in dorsal anterior cingulate sulcus reflect rewards but not travel time costs during foraging. Brain Neurosci Adv. 2019;

75. Porcelli AJ. An alternative to the traditional cold pressor test: the cold pressor arm wrap. J Vis Exp [Internet]. 2014;(83):e50849. Available from: http://www.ncbi.nlm.nih.gov/pubmed/24457998%5Cnhttp://www.pubmedcentral.nih.gov/articlerender.fcgi?artid=PMC4089407

76. Sokol-Hessner P, Raio CM, Gottesman SP, Lackovic SF, Phelps EA. Acute stress does not affect risky monetary decision-making. Neurobiol Stress. 2016;

77. Herrera AY, Nielsen SE, Mather M. Stress-induced increases in progesterone and cortisol in naturally cycling women. Neurobiol Stress. 2016;

78. Birkett MA. The Trier Social Stress Test Protocol for Inducing Psychological Stress. J Vis Exp. 2011;

79. McRae AL, Saladin ME, Brady KT, Upadhyaya H, Back SE, Timmerman MA. Strees reactivity: Biological and subjective responses to the cold pressor and Trier Social stressors. Hum Psychopharmacol. 2006;

80. Roemmich JN, Feda DM, Seelbinder AM, Lambiase MJ, Kala GK, Dorn J. Stress-induced cardiovascular reactivity and atherogenesis in adolescents. Atherosclerosis. 2011;

81. Allen MT, Matthews KA, Sherman FS. Cardiovascular reactivity to stress and left ventricular mass in youth. Hypertension. 1997;

82. Jaycox LH, Foa EB, Morral AR. Influence of emotional engagement and habituation on exposure therapy for PTSD. J Consult Clin Psychol. 1998;

83. Tsao J, Evans S, Seidman L, Lung, Zeltzer L, Naliboff B. Heart rate variability as a biomarker for autonomic nervous system response differences between children with chronic pain and healthy control children. J Pain Res. 2013;

84. Omurtag A, Abdel Baki SG, Chari G, Cracco RQ, Zehtabchi S, Fenton AA, et al. Technical and clinical analysis of microEEG: A miniature wireless EEG device designed to record high-quality EEG in the emergency department. Int J Emerg Med. 2012;

85. Delorme A, Makeig S. EEGLAB: an open source toolbox for analysis of single-trial EEG dynamics including independent component analysis. J Neurosci Methods. 2004;

