## Supplemental Table 1 for "A machine learning approach to identifying objective biomarkers of anxiety and stress"

| Demographics | Responses | Strong responders Mean (SD) |
|---|---|---|
| Total People in Each Group | 39 | 23 (59.0%) |
| Age | 39 | 26.94 (4.77) |
| Alcoholic Beverage Regularly (i.e. multiple times per week) | 38 | 27.60% |
| Coffee (Oz) consumed in the past week | 39 | 30.5 (41.89) |
| Current Hunger Level (1-10) | 38 | 3.73 (2.25) |
| Education (High School=0, College=1, Grad School=2) | 39 | 1.44 (0.51) |
| Energy Drink (Oz) consumed in the past week | 39 | 3.00 (9.30) |
| Hours of Sleep Last Night | 39 | 7.13 (1.31) |
| Oral Contraceptive Regularly (i.e. multiple times per week) | 22 | 20% |
| Recreational Drugs Regularly (i.e. multiple times per week) | 38 | 20.00% |
| Sex (Male=0, Female=1) | 39 | 62.50% |
| Soda (Oz) consumed in the past week | 39 | 12.75 (31.44) |
| Tea (Oz) consumed in the past week | 39 | 5.00 (12.77) |
| Tobacco Products Regularly (i.e. multiple times per week) | 39 | 6.30% |
| Weight (lbs) | 39 | 157.69 (34.05) |

| Weak /Non-responders Mean (SD) |
| --- |
| 16 (41.0%) |
| 23.91 (4.95) |
| 34.00% |
| 17.39 (23.09) |
| 4.04 (2.23) |
| 1.17 (0.39) |
| 0.00 (0.00) |
| 7.57 (0.84) |
| 42.20% |
| 0.00% |
| 52.20% |
| 3.74 (7.78) |
| 5.61 (11.29) |
| 4.30% |
| 157.61 (37.64) |
