## Supplemental Table 2 for "A machine learning approach to identifying objective biomarkers of anxiety and stress"

| Type | Split | Specificity | Sensitivity | Accuracy |
|---|---|---|---|---|
| Support Vector Machine | 9505 | 1 | 0.91 | 0.958 |
| Support Vector Machine | 9010 | 1 | 0.89 | 0.945 |
| Decision Tree Classifier | 8020 | 0.732484076 | 0.813253012 | 0.817 |
| K Nearest Neighbor | 8020 | 0.675159236 | 0.873493976 | 0.795 |
| Support Vector Machine | 8020 | 1 | 0.84939759 | 0.928 |
| Decision Tree Classifier | 7030 | 0.635036 | 0.841379 | 0.773 |
| K Nearest Neighbor | 7030 | 0.664234 | 0.834483 | 0.772 |
| Support Vector Machine | 7030 | 1 | 0.786207 | 0.896 |
| Decision Tree Classifier | 6040 | 0.70339 | 0.830645 | 0.718 |
| K Nearest Neighbor | 6040 | 0.686441 | 0.822581 | 0.693 |
| Support Vector Machine | 6040 | 1 | 0.782258 | 0.854 |
| Decision Tree Classifier | 5050 | 0.632653 | 0.769231 | 0.736 |
| K Nearest Neighbor | 5050 | 0.673469 | 0.759615 | 0.673 |
| Support Vector Machine | 5050 | 1 | 0.682692 | 0.825 |
| HRV | 5050 | 0.766667 | 0.804598 | 67 |
| HRV | 6040 | 0.805195 | 0.802083 | 69 |
| HRV | 7030 | 0.629412 | 0.85625 | 72 |
| HRV | 8020 | 0.615789 | 0.897849 | 74 |
| EEG | 5050 | 0.594059 | 0.755102 | 72 |
| EEG | 6040 | 0.643443 | 0.810345 | 72 |
| EEG | 7030 | 0.643885 | 0.863309 | 76 |
| EEG | 8020 | 0.708075 | 0.856688 | 79 |
| EDA | 5050 | 0.59633 | 0.80198 | 66 |
| EDA | 6040 | 0.627907 | 0.845528 | 68 |
| EDA | 7030 | 0.679739 | 0.865248 | 68 |
| EDA | 8020 | 0.670455 | 0.85 | 67 |
