## Supplemental Table 3 for "A machine learning approach to identifying objective biomarkers of anxiety and stress"

'FP1_T6_totalCoherencepct'
'HRV_Coherencepct'
'avgrrpct'
'hf_logpct'
'hfpct'
'lf_hf_ratio_logpct'
'pnn50pct'
'gammaPower_F8pct'
'deltaPower_O2pct'
'alphaPower_FCzpct'
'highbetaPower_F3pct'
'relative_gammaPower_C4pct'
'relative_alphaPower_FP2pct'
'relative_highbetaPower_FP2pct'
'relative_betaPower_F8pct'
'relative_alphaPower_T6pct'
'relative_thetaPower_FP2pct'
'relative_alphaPower_F8pct'
'relative_betaPower_C4pct'
'relative_deltaPower_T4pct'
'relative_gammaPower_F4pct'
'relative_deltaPower_T6pct'
'relative_thetaPower_C4pct'
'relative_deltaPower_T5pct'
'relative_lowbetaPower_T3pct'
'relative_alphaPower_F3pct'
'relative_alphaPower_O1pct'
'FP1_FP2_gammma_asymmetry_4pct'
'C3_C4_lowbeta_asymmetry_4pct'
'O1_O2_gammma_asymmetry_4pct'
'T5_T6_lowbeta_asymmetry_4pct'
'F7_F8_alpha_asymmetry_4pct'
'O1_O2_alpha_asymmetry_4pct'
'C3_C4_gammma_asymmetry_4pct'
'T5_T6_gammma_asymmetry_4pct'
'O1_O2_highbeta_asymmetry_4pct'
'T3_T4_highbeta_asymmetry_4pct'
'T3_T4_alpha_asymmetry_4pct'
'O1_O2_lowbeta_asymmetry_4pct'
'T3_T4_gammma_asymmetry_4pct'
'C3_C4_alpha_asymmetry_4pct'
'T3_T4_lowbeta_asymmetry_4pct'
'Engagement_Indexpct'
'FP1_FP2_lowbeta_asymmetry_4pct'
'F7_F8_lowbeta_asymmetry_4pct'
'T5_T6_highbeta_asymmetry_4pct'
'FP1_FP2_highbeta_asymmetry_4pct'

'F7_F8_highbeta_asymmetry_4pct'
'f3_f4_gammma_asymmetry_4pct'
'f3_f4_lowbeta_asymmetry_4pct'
'f3_f4_alpha_asymmetry_4pct'
'f3_f4_highbeta_asymmetry_4pct'
'HRpct'
'EDApct'
