## Supplemental Table 4 for "A machine learning approach to identifying objective biomarkers of anxiety and stress"

| algorithm | Split | Accuracy |
|---|---|---|
| Decision Tree Classifier | 8020 | 0.817 |
| Gradient Boosting Classifier | 8020 | 0.858 |
| K Nearest Neighbor | 8020 | 0.795 |
| Logistic Regression | 8020 | 0.713 |
| Naive Bayes Classifier | 8020 | 0.629 |
| Random Forest Classifier | 8020 | 0.858 |
| Support Vector Machine | 8020 | 0.928 |
| Decision Tree Classifier | 7030 | 0.773 |
| Gradient Boosting Classifier | 7030 | 0.833 |
| K Nearest Neighbor | 7030 | 0.772 |
| Logistic Regression | 7030 | 0.727 |
| Naive Bayes Classifier | 7030 | 0.627 |
| Random Forest Classifier | 7030 | 0.819 |
| Support Vector Machine | 7030 | 0.896 |
| Decision Tree Classifier | 6040 | 0.718 |
| Gradient Boosting Classifier | 6040 | 0.796 |
| K Nearest Neighbor | 6040 | 0.693 |
| Logistic Regression | 6040 | 0.686 |
| Naive Bayes Classifier | 6040 | 0.591 |
| Random Forest Classifier | 6040 | 0.771 |
| Support Vector Machine | 6040 | 0.854 |
| Decision Tree Classifier | 5050 | 0.736 |
| Gradient Boosting Classifier | 5050 | 0.773 |
| K Nearest Neighbor | 5050 | 0.673 |
| Logistic Regression | 5050 | 0.747 |
| Naive Bayes Classifier | 5050 | 0.575 |
| Random Forest Classifier | 5050 | 0.782 |
| Support Vector Machine | 5050 | 0.825 |
