## Supplemental Table 5 for "A machine learning approach to identifying objective biomarkers of anxiety and stress"

| Trait Anxiety | 2-state: low high (35-thresh) | | | |
|---|---|---|---|---|
| Classifier | Features | Accuracy | | |
| | | 90-10 | 95-5 | LOOCV |
| complex trees | $SUDS_{T}$+$STAI_{B+T}$+ $SC_{B+T}$ | 73 | 81.1 | 83.8 |
| boosted trees | $SUDS_{T}$+$STAI_{B+T}$+ $SC_{B+T}$ | 70.3 | 78.4 | 73 |
| weighted KNN (10) | $SUDS_{T}$+$STAI_{B+T}$+ $SC_{B+T}$ | 81.1 | 70.3 | 73 |
| Logistic regression | $SUDS_{T}$+$STAI_{B+T}$+ $SC_{B+T}$ | 56.8 | 62.2 | 59.5 |
| LDA | $SUDS_{T}$+$STAI_{B+T}$+ $SC_{B+T}$ | 62.2 | 62.2 | 62.2 |
